# Reversion of Ebolavirus Disease from a Single Intramuscular Injection of a pan-Ebolavirus Immunotherapeutic

**DOI:** 10.1101/2022.01.06.475142

**Authors:** Erin Kuang, Robert W. Cross, Maria McCavitt-Malvido, Dafna M. Abelson, Viktoriya Borisevich, Krystle N. Agans, Neil Mlakar, Arumugapradeep Marimuthu, Daniel J. Deer, William S. Shestowsky, Do Kim, Joan B. Geisbert, Larry Zeitlin, Crystal L. Moyer, Chad J. Roy, Thomas Geisbert, Zachary A. Bornholdt

**Affiliations:** Mapp Biopharmaceutical, Inc., San Diego, CA 92121, USA; Galveston National Laboratory, University of Texas Medical Branch, Galveston, TX 77555, USA; Department of Microbiology and Immunology, University of Texas Medical Branch, Galveston, TX 77555, USA; Division of Microbiology, Tulane National Primate Research Center, Covington, LA 70447, USA; Department of Microbiology and Immunology, Tulane School of Medicine, New Orleans, LA 70112, USA

**Keywords:** Monoclonal antibody, ebolavirus, intravenous infusion, intramuscular injection, half-life extension, post-exposure prophylaxis, therapeutic

## Abstract

Intravenous administration (IV) of antiviral monoclonal antibodies (mAbs) is challenging due to limited resources for performing infusions during an ongoing epidemic. An ebolavirus therapeutic administered via intramuscular (IM) injection would reduce these burdens and allow rapid treatment of exposed individuals during an outbreak. Here, we demonstrate how MBP134, a two mAb pan-ebolavirus cocktail, reverses the course of *Sudan ebolavirus* (SUDV/Gulu) disease with a single IV or IM dose in non-human primates (NHPs) as far as five days post-exposure. Furthermore, we investigated the utility of adding half-life extension mutations to the MBP134 mAbs, ultimately creating a half-life extended cocktail designated MBP431. MBP431 demonstrated an extended serum half-life *in vivo* and offered complete or significant protection with a single IM dose delivered as a post-exposure prophylactic (PEP) or therapeutic in NHPs challenged with EBOV. These results support the use of MBP431 as a rapidly deployable IM medical countermeasure against every known ebolavirus.

## 1.1 INTRODUCTION

Ebolaviruses are members of the family *Filoviridae* known to cause severe hemorrhagic fever in both humans and nonhuman primates (NHPs), with human mortality rates in some outbreaks approaching 90% (Burk et al., 2016). There are now six known ebolavirus species: *Zaire ebolavirus* (EBOV), *Sudan ebolavirus* (SUDV), *Bundibugyo ebolavirus* (BDBV), *Taï forest ebolavirus* (TAFV), *Reston ebolavirus* (RESTV) and *Bombali ebolavirus* (BOMV) (Burk et al., 2016; Goldstein et al., 2018). While two monoclonal antibody (mAb) products, REGN-EB3 (Atoltivimab, Maftivimab, and Odesivimab) and mAb114 (Ansuvimab), have been FDA approved for the treatment of EBOV infection, they are not active against the remaining five ebolavirus species (Iversen et al., 2020). In contrast, the two mAbs, ADI-15878 and ADI-23774, that make up the MBP134 cocktail, target highly conserved epitopes on the ebolavirus glycoprotein (GP) spike and inhibit GP-mediated membrane fusion across every known species of ebolavirus (Bornholdt et al., 2019; Bornholdt et al., 2016; Murin et al., 2018; Wec et al., 2019; Wec et al., 2017). We previously demonstrated MBP134 could reverse the course of ebolavirus disease (EVD) in NHPs challenged with EBOV, SUDV, or BDBV with a single intravenous (IV) dose (Bornholdt et al., 2019). However, IV administration of a therapeutic in an EBOV treatment unit (ETU) is resource intensive, time consuming, and requires medical personnel to deliver the drug within a clinical setting.

A single dose IM pan-ebolavirus PEP/therapeutic would dramatically increase the number of exposed individuals who could be rapidly treated in settings with limited medical resources and reduce infection risk for medical professionals in ETUs. The high dosing (≥50 mg/kg) required for protective efficacy of currently approved EBOV therapeutics necessitates IV administration, limiting rapid large-scale deployment (Mulangu et al., 2019). Additionally, by targeting epitopes limited in activity to only EBOV, both REGN-EB3 and mAb114 do not address public health threats posed by emerging divergent ebolavirus species (e.g. SUDV or BDBV), a significant liability that arose with SARS-CoV-2 immunotherapeutics which has led to reduced levels of protection over time (Planas et al., 2021). Thus, we set out to determine if MBP134, a highly potent pan-ebolavirus immunotherapeutic, could be administered at lower IV or IM doses. We also examined the impact serum half-life extensions would have on the MBP134 mAbs to potentially increase their clinical utility for use in prophylaxis or PEP scenarios.

## 1.2 MATERIALS AND METHODS

### Study design

The objective of the research performed here was to determine if a two antibody pan-ebolavirus cocktail administered intramuscularly could provide protective efficacy in non-human primates challenged with either SUDV or EBOV, potentially altering ebolavirus outbreak response paradigms in the clinic. For the first study, nine healthy, adult rhesus macaques (*Macaca mulatta*) of Chinese origin (PreLabs) ranging in age from ∼4-5 years and weighing 5.4-6.3 kg were challenged intramuscularly (IM) in the left quadricep with a 1000 PFU target dose of SUDV (Gulu variant). Treatment was initiated 5 days after infection as a single IV dose of MBP134. The duration of this study was 28 days. For the second study, eleven healthy, adult rhesus macaques of Chinese origin (Prelabs) ranging in age from ∼ 4-6 years and weighing 4.0-6.6 kg were challenged IM in the left quadricep with a 1000 PFU target dose of SUDV (Gulu variant, SUDV/Gulu). Treatment was initiated 3 or 5 days after infection as a single IM dose of MBP134. In order to maintain blinding of the groups, every animal received an IM injection of either MBP134 or phosphate buffered saline (PBS) on D3 and D5 (control received two injections of PBS) from deidentified vials that were randomly assigned prior to challenge. The duration of this study was 28 days. For the third study, eleven healthy, adult rhesus macaques of Chinese origin (Prelabs) ranging in age from ∼ 3-4 years and weighing 4.2-5.2 kg were challenged IM in the left quadricep with a 1000 PFU target dose of EBOV (Kikwit variant). Treatment was initiated 3 days after infection as a single IM dose of MBP431. The duration of this study was 28 days. For the fourth study, six healthy, adult rhesus macaques of Chinese origin (Prelabs) ranging in age from ∼ 3-4 years and weighing 4.0-5.2 kg were challenged IM in the left quadricep with a 1000 PFU target dose of EBOV (Kikwit variant). Treatment was initiated 4 days after infection as a single IM dose of MBP431. The duration of this study was 28 days. For all four NHP studies, assignment to each treatment group or control was determined prior to challenge by randomization with effort made to maintain a balanced sex ratio. All studies were blinded to the research staff. The macaques were monitored daily and scored for disease progression with an internal filovirus humane endpoint scoring sheet approved by the UTMB Institutional Animal Care and Use Committee (IACUC). The scoring changes measured from baseline included posture and activity level, attitude and behavior, food intake, respiration, and disease manifestations, such as visible rash, hemorrhage, ecchymosis, or flushed skin. A score of ≥ 9 indicated that an animal met the criteria for euthanasia.

### Expression, purification, and formulation of monoclonal antibodies from an engineered CHOK1-AF cell line

CHOK1-AF cells stably expressing the ADI-23774 and ADI-15878 mAbs were generated as previously described (Bornholdt et al., 2019). Briefly, a dual plasmid system containing expression cassettes for the heavy and light chains of the target mAbs were co-transfected by random integration via chemical means into a modified CHOK1 host cell line (CHOK1-AF) which yields afucosylated glycans on expressed mAbs. Stable selection was initiated with the inclusion of MSX as a selection agent 24 hours post transfection. Upon completion of the final expansion, the culture was maintained in fed-batch for 14 days, after which the supernatant was clarified via filtration and subsequently sterile filtered (0.2 μm) into a 20 L bioprocess bag (Thermo Fisher) prior to protein A purification. For the first NHP SUDV/Gulu challenge study MBP134 was formulated for IV delivery in 20 mM sodium citrate, 10 mM glycine, 8% sucrose, 0.01% polysorbate 80 (PS80), pH 5.5 (Buffer 1) at 7.3 mg/mL (low dose treatment) or 24.1 mg/mL (high dose treatment). In order to support IM delivery of MBP134 in the second NHP SUDV/Gulu challenge study, the drug product was formulated in Buffer 1 at 60.7 mg/mL. MBP431 was first formulated in Buffer 1 at 20.4 mg/mL (low dose) and 60.8 mg/mL (high dose) for the first EBOV/Kikwit challenge study and later formulated in 10 mM histidine, 5% sorbitol, 0.02% PS80, pH 6.0 at 22.4 mg/mL for the final EBOV/Kikwit challenge study.

### Generation of anti-idiotype antibodies

The anti-idiotype antibodies targeting either ADI-15878 or ADI-23774 were raised by Bio-Rad using the HuCal PLATINUM synthetic Fab library (Prassler et al., 2011). Anti-idiotype Fabs were screened for binding against the target Fab, IgG1, and against a negative IgG1 isotype control antibody yielding the final candidate anti-idiotypes for ADI-15878 and ADI-23774, anti-idiotype-878 (AI878) and anti-idiotype-774 (AI774), respectively. The sequences for each anti-idiotype Fab were provided from Bio-Rad. The Fab heavy and light chains of AI774 and AI878 were individually cloned into a pcDNA3 vector (Invitrogen). A 6x-His tag was added to the C-terminus of each Fab heavy chain sequence. Plasmids containing the Fab heavy chains and light chains were chemically co-transfected into an afucosylated ExpiCHO cell line using the ExpiFectamine CHO Transfection Kit (Gibco). After 9 days of transient expression, the supernatant containing either the AI774 or AI878 Fab was clarified via centrifugation and 0.22 µm filtered prior to purification. The Fabs were purified by affinity chromatography on a 5 mL HisTrap FF crude column (Cytiva). Following loading, the column was washed with PBS + 5 mM imidazole, and the Fabs were eluted with PBS + 250 mM imidazole. For AI774, the nickel column eluate was further purified on a 1 mL HiTrap LambdaFabSelect column (Cytiva). The column was washed with PBS and the AI774 Fab was eluted with IgG Elution Buffer (Pierce). Prior to storage at -70°C, the AI774 Fab was neutralized to pH 7 with 1 M Tris, pH 8.0.

### Pharmacokinetics study in rhesus macaques

A PK study in rhesus macaques was used to determine the serum half-life of the four Fc-engineered MBP134 mAbs to assess their potential as an immunoprophylactic countermeasure. Two NHPs received ADI-23774^YTE^ and ADI-15878^YTE^ (MBP431 cocktail), while three NHPs received ADI-23774^XTND^ and ADI-15878^XTND^ (MBP432 cocktail). Each NHP was given an intravenous dose of 14 mg/kg of the assigned set of mAbs. Blood samples were obtained from the NHPs prior to treatment; 5 hours post-treatment; and 1, 2, 4, 7, 10, 16, 21, 78, 92, 136, and 150 days post-treatment. The concentration of each mAb in each serum sample was quantified longitudinally by enzyme linked immunosorbent assay (ELISA). 96-well half-area ELISA microplates (Greiner Bio-One) were coated with 2 µg/ml of mAb-specific anti-idiotype Fab (AI87 or AI47) at 4 °C overnight. The next morning, plates were blocked with 100 µl/well of SuperBlock (Thermo Scientific) at room temperature for 30 minutes. Serum samples were diluted 1:5000 in 0.05% Tween-20 PBS (PBST) with 1% BSA, and then added to the blocked wells at room temperature for 1 hour. Similarly, 12-point standard curves were generated for each mAb by adding 1/3-fold serial dilutions (starting at 3 µg/ml for ADI-23774 and 10 µg/ml for ADI-15878) to the blocked wells. Plates were washed 3 times with PBST, and then incubated with a 1:10000 dilution (in PBST with 1% BSA) of goat anti-human Fc HRP-conjugated antibody (Invitrogen) at room temperature for 1 hour. Following 3 washes with PBST, plates were developed by addition of 3,3’, 5,5’-tetramethylbenzidine substrate (KPL SureBlue, SeraCare) and stopped by addition of NH_2_SO_4_. Absorbance values were measured at 450 nm using the Perkin Elmer EnVision multimode plate reader.

### Pharmacokinetic data analysis

Data obtained from the PK study was analyzed in Prism 8 (GraphPad). Based on separate standard curves generated for each mAb, the serum concentration of each mAb for each timepoint was quantified and plotted against the number of days post-treatment. The serum half-life of each mAb was calculated using nonlinear regression, in which a two-phase exponential decay model (with y constrained to plateau at 0) was used to fit the data. The value reported here is the terminal elimination half-life.

### University of Texas Medical Branch (UTMB) ethics statement

Animal studies were performed in Biosafety level (BSL)-4 biocontainment at the University of Texas Medical Branch (UTMB) and approved by the UTMB Institutional Biosafety Committee (IBC) and IACUC. Animal research was conducted in compliance with UTMB IACUC, Animal Welfare Act, and other federal statutes and regulations relating to animals. The UTMB animal research facility is fully accredited by the Association for Assessment and Accreditation of Laboratory Animal Care and adhere to principles specified in the eighth edition of the Guide for the Care and Use of Laboratory Animals, National Research Council.

### Tulane National Primate Research Center (TNPRC) ethics statement

The Tulane University Institutional Animal Care and Use Committee approved all procedures used during this study. The Tulane National Primate Research Center (TNPRC) is accredited by the Association for the Assessment and Accreditation of Laboratory Animal Care (AAALAC no.000594). The U.S. National Institutes of Health (NIH) Office of Laboratory Animal Welfare number for TNPRC is A3071-01.

### Challenge viruses

*Zaire ebolavirus* (EBOV) isolate 199510621 (strain Kikwit) originated from a 65-year-old female patient who had died on 5 May 1995. The study challenge material was from the second Vero E6 passage of EBOV isolate 199510621. Briefly, the first passage at UTMB consisted of inoculating CDC 807223 (passage 1 of EBOV isolate 199510621) at an MOI of 0.001 onto Vero E6 cells (ATCC CRL-1586). The cell culture fluids were subsequently harvested at day 10 post infection and stored at -80°C as ∼1 ml aliquots. Deep sequencing indicated the EBOV was greater than 98% 7U (consecutive stretch of 7 uridines). No detectable mycoplasma or endotoxin levels were measured (□ 0.5 endotoxin units (EU)/ml). *Sudan ebolavirus* (SUDV) isolate 200011676 (strain Gulu) originated from a 35-year-old male patient who had died on 16 October 2000. The study challenge material was from the second Vero E6 cell passage of SUDV isolate 200011676. Briefly, the first passage at UTMB consisted of inoculating CDC 808892 (CDC passage 1 of SUDV isolate 200011676) at an MOI of 0.001 onto Vero E6 cells (ATCC CRL-1586). The cell supernatants were subsequently harvested at day 7 post infection and stored at - 80°C as ∼ 1 ml aliquots. No detectable mycoplasma or endotoxin levels were measured (□ 0.5 EU/ml). Genomic sequencing of the resulting seed revealed a 92.33% 7U genotype prevailance at the glycoprotein editing site (Geisbert et al., 2015).

### Hematology and serum biochemistry

Total white blood cell counts, white blood cell differentials, red blood cell counts, platelet counts, hematocrit values, total hemoglobin concentrations, mean cell volumes, mean corpuscular volumes, and mean corpuscular hemoglobin concentrations were analyzed from blood collected in tubes containing EDTA using a Vetscan HM5 laser based hematologic analyzer (Zoetis). Serum samples were tested for concentrations of albumin, amylase, alanine aminotransferase (ALT), aspartate aminotransferase (AST), alkaline phosphatase (ALP), blood urea nitrogen (BUN), calcium, creatinine (CRE), C-reactive protein (CRP), gamma-glutamyltransferase (GGT), glucose, total protein, and uric acid by using a Piccolo point-of-care analyzer and Biochemistry Panel Plus analyzer discs (Abaxis).

### RNA isolation from SUDV- and EBOV-infected macaques

On procedure days, 100 μl of blood from K2-EDTA collection tubes was collected prior to centrifugation and was added to 600 μl of AVL viral lysis buffer with 6 μL carrier RNA (Qiagen) for RNA extraction. For tissues, approximately 100 mg was stored in 1 ml RNAlater (Qiagen) for at least 4 days for stabilization. RNAlater was completely removed, and tissues were homogenized in 600 μl RLT buffer and 1% betamercaptoethanol (Qiagen) in a 2 mL cryovial using a tissue lyser (Qiagen) and 0.2mm ceramic beads. The tissues sampled included axillary and inguinal lymph nodes, liver, spleen, kidney, adrenal gland, lung, pancreas, urinary bladder, ovary or testis, and eye. All blood samples were inactivated in AVL viral lysis buffer, and tissue samples were homogenized and inactivated in RLT buffer prior to removal from the BSL-4 laboratory. Subsequently, RNA was isolated from blood using the QIAamp viral RNA kit (Qiagen), and from tissues using the RNeasy minikit (Qiagen) according to the manufacturer’s instructions supplied with each kit.

### Quantification of viral load

Primers and probe targeting the VP30 gene of EBOV and the L gene of SUDV were used for real-time quantitative PCR (RT-qPCR) with the following probes: EBOV, 6-carboxyfluorescein (FAM)-5= CCG TCA ATC AAG GAG CGC CTC 3=-6 carboxytetramethylrhodamine (TAMRA); SUDV, FAM-5=CAT CCA ATC AAA GAC ATT GCG A 3=-TAMRA (Life Technologies).Viral RNA was detected using the CFX96 detection system (Bio-Rad Laboratories, Hercules, CA) in one-step probe RT-qPCR kits (Qiagen) with the following cycle conditions: EBOV, 50°C for 10 min, 95°C for 10 s, and 40 cycles of 95°C for 10 s and 57°C for 30 s; SUDV, 50°C for 10 min, 95°C for 10 s, and 40 cycles of 95°C for 10 s and 59°C for 30 s. Threshold cycle (CT) values representing viral genomes were analyzed with CFX Manager software, and the data are shown as genome equivalents (GEq). To create the GEq standard, RNA from viral stocks was extracted, and the number of strain-specific genomes was calculated using Avogadro’s number and the molecular weight of each viral genome. Virus titration was performed by plaque assay using Vero E6 cells (ATCC CRL-1586) from all plasma samples as previously described (Bornholdt et al., 2019). Briefly, increasing 10-fold dilutions of the samples were adsorbed to Vero E6 cell monolayers in duplicate wells (200 μl) and overlaid with 0.8% agarose in 1x Eagles minimum essentials medium (MEM) with 5% FBS and 1% P/S. After 6 to 12 days incubation at 37°C/5% CO_2_, neutral red stain was added and plaques were counted after 48 hour incubation. The limit of detection for this assay was 5 PFU/mL.

## 1.3 RESULTS

### 1.3.1 A single IV dose of MBP134 protects NHPs from lethal SUDV challenge

Previously, we demonstrated MBP134 protects NHPs challenged with SUDV/Boniface and ferrets challenged with SUDV/Gulu (Bornholdt et al., 2019). Here, MBP134 was further evaluated for therapeutic efficacy in a fully blinded study in the lethal SUDV/Gulu rhesus macaque model. Nine animals were challenged IM with a target dose of 1000 plaque forming units (PFU) of SUDV/Gulu (actual dose = 1013 PFU). One animal was assigned to the untreated control group with historical control data leveraged to support statistical analyses. The treatment groups (n=4/group) received either a single 25 mg/kg or a single 7.5 mg/kg IV dose of MBP134 on day 5 (D5) post-infection (PI) (**Figure 1A**). Both the 25 mg/kg and 7.5 mg/kg dose of MBP134 reversed the course of SUDV/Gulu disease and protected all of the challenged animals, while the control animal succumbed to SUDV/Gulu infection on D8 PI (**Figure 1A**). Historical controls receiving the same viral inoculum under identical protocols had a mean time to death of 9 days PI (n=5). Retrospectively, all of the animals were confirmed to be qRT-PCR positive for SUDV/Gulu infection at time of treatment with viremia ranging from 5.7 to 10.5 log_10_ genomic equivalents (GEQ)/mL (**Figure 1B**). Infectious SUDV/Gulu virus was found in the blood at greater than 4.5 log_10_ PFU/mL on D5 PI in all but two animals. NHP-4 showed 2.7 Log_10_ PFU/mL and NHP-7 tested just above the lower limit of detection (LLOD) at 1.4 Log_10_ PFU/mL on D5 PI (**Figure 1C**). MBP134-treated animals were PCR negative, or below the LLOD, nine days post-treatment and had no detectable infectious virus (PFUs) by the first post-treatment collection point on D8 PI (**Figure 1B and 1C**). All of the NHPs showed clinical signs of SUDV disease as reflected by the presence of fever, reduced platelet count, lymphopenia, elevated C-reactive protein (CRP) levels, and clinical scoring prior to treatment on D5 PI, supportive of a therapeutic indication (**Figure 1D-H**). Of note, both the control and the 25 mg/kg treatment group displayed elevated levels of alanine aminotransferase (ALT) by D8 PI, returning to baseline levels by D14 PI for the treated animals. In contrast, ALT levels of the 7.5 mg/kg treatment group remained relatively unchanged from baseline (**Figure 1I**). However, in both treatment groups aspartate aminotransferase (AST) and alkaline phosphatase (ALP) remained at baseline levels, demonstrating MBP134 inhibited the progression of liver damage typical of SUDV infection as observed in the control (**Figure S1**). These data demonstrate the single dose therapeutic potency of MBP134 for the treatment of SUDV/Gulu infection.

**Figure 1.**
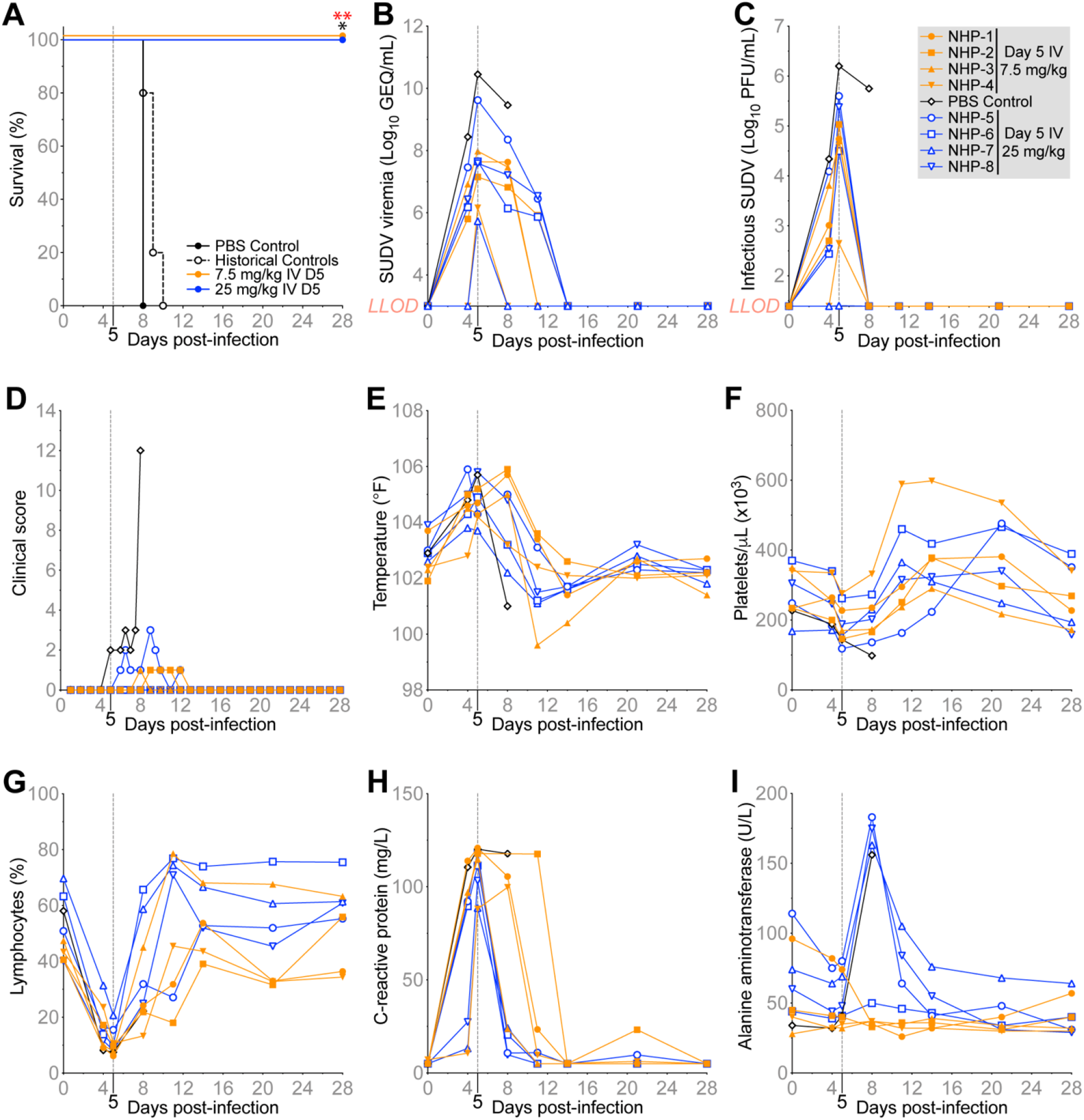
Evaluation of MBP134 delivered IV in NHPs challenged with SUDV/Gulu. **(A)** Survival curves for NHPs challenged IV with 1000 PFU of SUDV/Gulu. Animals received either PBS (black), a 25 mg/kg dose (blue), or a 7.5 mg/kg dose (orange) of MBP134 on D5 PI. P values were determined using a Log-rank Mantel-Cox test, against the in-study control (* = 0.0455) alone or the in-study control combined with the historical controls (** = 0.0047, shown in red). **(B)** Average GEQ/mL of SUDV/Gulu and **(C)** infectious SUDV/Gulu (PFU/mL) in the blood of each NHP. **(D)** Clinical scores, **(E)** body temperatures, **(F)** platelet counts, **(G)** lymphocyte levels, **(H)** C-reactive protein (CRP) levels, and **(I)** alanine aminotransferase (ALT) levels of each NHP. Legend for panels **(B-I)** appears in top right-hand corner (grey box), *LLOD* = lower limit of detection (3 log_10_ GEQ/mL and 1.39 PFU/mL). See also Figure S1.

### 1.3.2 MBP134 maintains therapeutic efficacy when delivered as an IM injection

The complete therapeutic protection observed in NHPs treated with a single 7.5 mg/kg IV dose of MBP134 suggested MBP134 may be sufficiently potent to support administration via IM injection. To evaluate MBP134 as either an IM injectable PEP or therapeutic drug, rhesus macaques were treated with a two-site IM 15 mg/kg total dose of MBP134 on either on D3 PI (NHP 1-5) or D5 PI (NHP 6-10). One animal was assigned to the untreated control group, with historical control data leveraged to support statistics (n=1 from the previous study and n=5 from previously published studies (Thi et al., 2016)). All NHPs were challenged IM with a target dose of 1000 PFU of SUDV/Gulu (actual dose = 888 PFU) using the same viral stock as historical studies (Thi et al., 2016). MBP134, when administered as an IM PEP injection on D3 PI, either prevented (NHP 1-3) or reversed (NHP 4-5) the course of SUDV/Gulu disease, with all animals surviving. Similarly, IM treatment D5 PI also provided significant levels of protection (80%; p-value = 0.0181) (**Figure 2A**). Unexpectedly, while the untreated control animal did become viremic and displayed a prolonged course of disease compared to the treated animals (**Figure 2**), the animal survived and represents the first untreated NHP (of seven total controls leading up to this study) to have survived a challenge from this SUDV/Gulu viral stock. As anticipated, most of the animals treated D3 PI (NHP 1-3) had no detectable infectious virus and remained RT-PCR negative throughout the study. However, NHP-4 and NHP-5 showed signs of an acute course of SUDV infection with 5.1 and 6.0 log_10_ SUDV/Gulu GEQ/mL detected on D3 PI, respectively. Infectious virus was found in NHP-5 on D3 and immediately dropped below the LLOD post-treatment. In contrast, all five animals (NHP 6-10) in the D5 treatment group were RT-PCR positive prior to treatment, ranging from 6.7 to 10.0 log_10_ SUDV/Gulu GEQ/mL (**Figure 2B**), and displayed high titers of infectious virus at time of treatment, ranging from 3.9 to 6.7 Log_10_ PFU/mL, similar to our first study here and previous SUDV challenge studies (Thi et al., 2016). Consistent with the previous experiment, no infectious virus was detectable following MBP134 IM treatment by the next sample collection point on D8 PI (**Figure 2C**). Prior to treatment, the D5 treatment group registered onset of fever and thrombocytopenia, while most NHPs treated on D3 PI avoided these clinical signs (**Figure 2E-F**). All animals developed lymphopenia and elevated CRP at time of treatment (**Figure 2G-H**). Of significance, NHP-8, the only treated animal to succumb to infection, displayed an acute course of SUDV disease compared to the other animals in the study (**Figure 2D**). NHP-8 had the highest level of viremia at time of treatment and showed signs of a severe systemic SUDV infection with significant levels of virus (GEQ per gram) in its lungs, adrenal glands, and kidneys upon necropsy (**Figure S2**). Additionally, this animal had the highest GEQ/mL and PFU/mL in blood of the cohort prior to treatment on D5 PI (**Figure 2B and 2C**). Notably, elevated AST levels appeared much earlier in NHP-8 than in the rest of the cohort, likely impacting chances of recovery (**Figure 2I**). The data here demonstrate MBP134 can prevent or reverse the course of SUDV/Gulu disease when administered as a PEP or as a therapeutic via IM injection.

**Figure 2.**
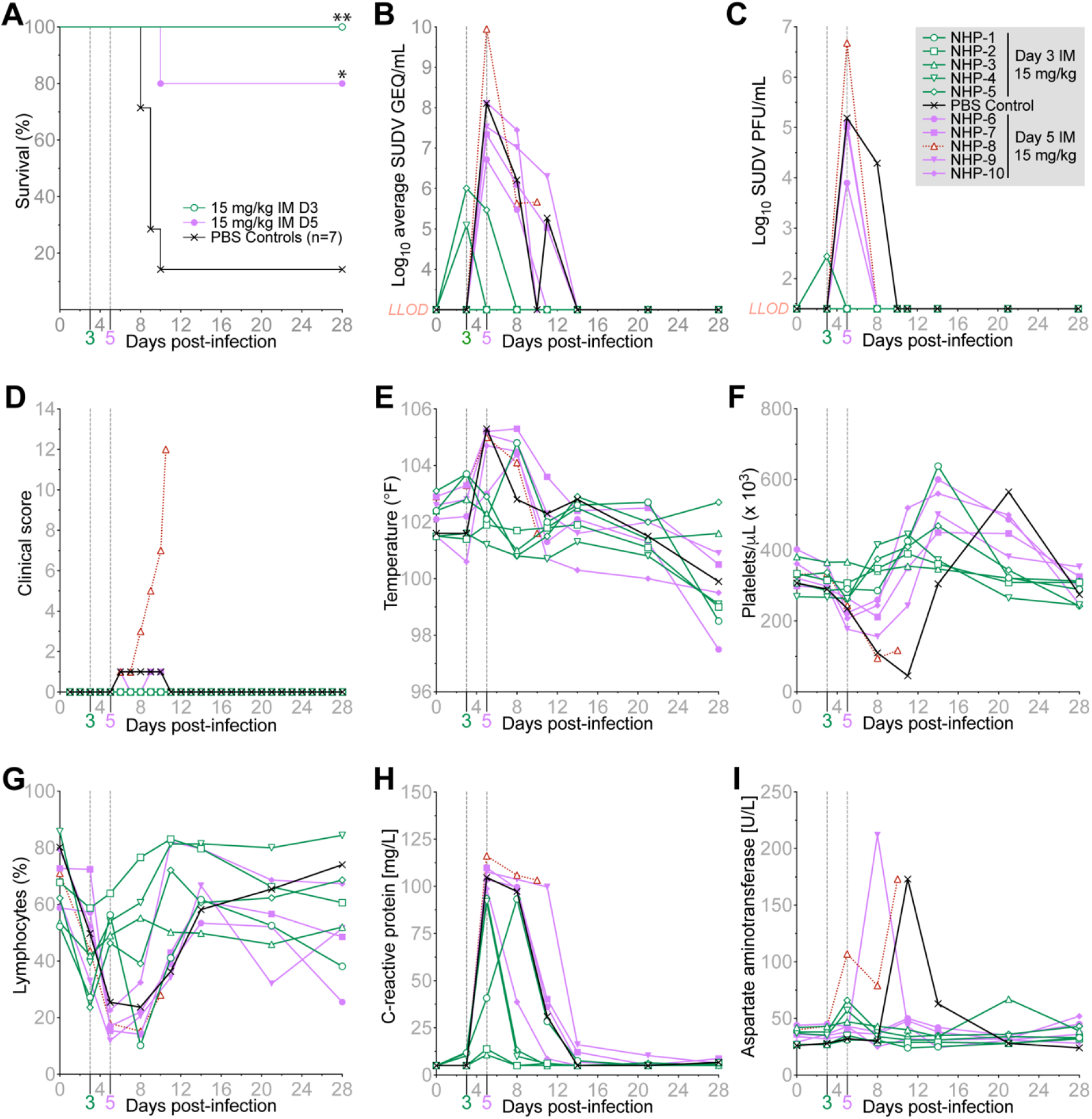
Evaluation of MBP134 as an IM injectable PEP or therapeutic drug in NHPs challenged with SUDV/Gulu. **(A)** Survival curves for NHPs challenged IM with 1000 PFU of SUDV/Gulu. Animals received either a 15 mg/kg dose of MBP134 on D3 PI (green) or D5 PI (purple). Historical controls and the in-study control are shown in black. P values were determined using a Log-rank Mantel-Cox test, against the combined controls with P values of 0.006 (**) and 0.0181(*) for the D3 and D5 treated animals, respectively. **(B)** Average GEQ/mL of SUDV/Gulu in the blood of each animal. NHP-8, the only treated animal that succumbed to infection, is shown in red. **(C)** Infectious SUDV/Gulu (PFU/mL) in the blood of each NHP. **(D)** Clinical scores, **(E)** body temperatures, **(F)** platelet counts, **(G)** lymphocyte levels, **(H)** C-reactive protein (CRP) levels, and **(I)** aspartate aminotransferase (AST) levels of each NHP. Legend for panels **(B-I)** appears in top right-hand corner (grey box), *LLOD* = lower limit of detection (3 log_10_ GEQ/mL and 1.39 PFU/mL). See also Figure S2.

### 1.3.3 Half-life extension mutations expand the clinical utility of MBP134

Previous studies have shown that YTE mutations (M252Y/S254T/T256E) and LS mutations (M428L/N434S) incorporated into an IgG Fc region can significantly extend antibody serum half-life by promoting pH-dependent recycling via binding to human FcRn (Dall’Acqua et al., 2006; Zalevsky et al., 2010). Studies also suggest these mutations can contribute to increased efficacy by maintaining higher levels of circulating immunotherapies for longer periods of time (Ko et al., 2014). Here, the YTE or LS mutations were introduced into the Fc region of ADI-15878 and ADI-23774, creating two half-life extended cocktail variants designated MBP431 and MBP432, respectively. A pharmacokinetics study was conducted in rhesus macaques to determine whether the YTE or the LS mutations were optimal for prolonging MBP134 serum availability. NHPs were given a 14 mg/kg IV dose of either MBP431 (n=2) or MBP432 (n=3). The serum concentration of each antibody was quantified using an anti-idiotype mAb specific to either ADI-15878 or ADI-23774 (**Figure 3A and 3B**). The calculated serum half-life of ADI-15878^YTE^ was 44 days, while that of ADI-15878^LS^ was 29 days (**Figure 3C**). The serum concentration of ADI-15878^LS^ began to drop off at a faster rate than ADI-15878^YTE^ around 50 days post-administration. The half-life of ADI-23774^YTE^ was 34.64 days, while that of ADI-23774^LS^ was reduced to 7.1 days (**Figure 3D**). The serum concentration of ADI-23774^LS^ dropped off dramatically after about 25 days post-administration at a comparatively faster rate than ADI-23774^YTE^. Of note, human clinical studies with human mAbs containing YTE mutations have reported half-lives of 80-112 days (Saunders, 2019). Thus, MBP431 was selected over MBP432 for further evaluation in NHP challenge studies due to its potential to provide months of protection.

**Figure 3.**
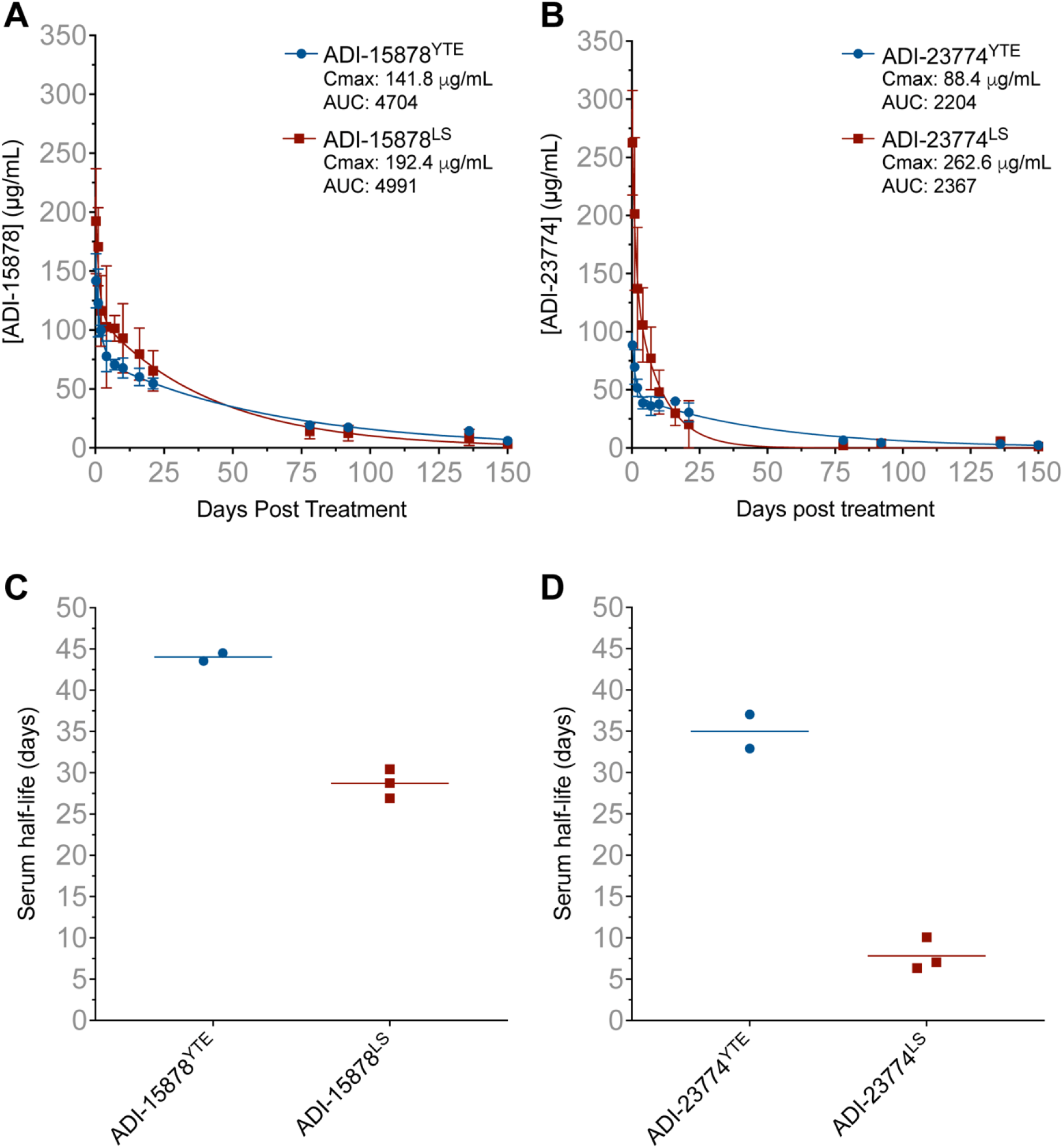
Pharmacokinetic analyses of half-life extension variants of MBP134 in NHPs. **(A)** Serum concentrations of ADI-15878^YTE^ (blue) and ADI-15878^LS^ (red) and **(B)** serum concentrations of ADI-23774^YTE^ (blue) and ADI-23774^LS^ (red) at each collection time point. The maximal concentration (Cmax) and area under the curve (AUC) are displayed in the legends for panels **(A-B)**. Errors bars represent ± standard deviation from the mean. Panels **(C-D)** display the calculated serum half-life for each mAb variant in each individual animal.

### 1.3.4 MBP431 provides NHPs significant PEP and therapeutic protection from EBOV

Building off of the efficacy demonstrated by MBP134 when delivered IM to NHPs challenged with SUDV, we evaluated the efficacy of MBP431 in NHPs challenged with EBOV/Kikwit. Eleven NHPs were challenged IM with a target dose of 1000 PFU of EBOV/Kikwit (actual dose = 1088 PFU). To evaluate MBP431 as PEP, the treatment groups (n=5/group) received a two-site IM injection of MBP431 at either a 15 mg/kg or 5 mg/kg total dose on D3 PI. One control animal received PBS. Both dosing regimens provided complete protection in contrast to the PBS treated control (p-value = 0.0003), which met euthanasia criteria on D7 PI (**Figure 4A**). Historic controls inoculated with the same viral lot under identical protocols had a mean time to death of 7 days PI (n = 12). Animals in both groups showed signs of active EBOV infection on D3 PI prior to treatment with virus detected via qRT-PCR from the LLOD to 6.8 log_10_ GEQ/mL. Coinciding with the RT-PCR data, animals displayed levels of circulating infectious virus ranging from the LLOD to 3.4 log_10_ PFU/mL (**Figure 4B and 4C**). However, either dose delivered on D3 PI inhibited significant advancement of EBOV infection, with viremia levels remaining below the LLOD throughout the remainder of the study. Based on these results, the time to treat was delayed by 24 hours to D4 PI in a follow-up study in which six NHPs were challenged with EBOV (target dose 1000 PFU, actual dose 1225 PFU). One control animal received PBS, while the other five received a two site IM injection of MBP431 for a 5 mg/kg total dose. MBP431 provided significant, but reduced levels of protection, with 60% of treated NHPs surviving (**Figure 4D**). All of the animals in the study were qRT-PCR positive prior to treatment on D4 PI. Of significance, survivors (NHP-1, NHP-4, and NHP-5) were grouped around 7-8 log_10_ GEQ/mL, while non-survivors (NHP-2, NHP-3, and the control) were grouped at ∼10 log_10_ GEQ/mL on the day of treatment (**Figure 4E**). Similarly, infectious virus was present in blood samples from all the animals prior to treatment on D4 PI, with survivors having viral loads up to 3.1 log_10_ PFU/mL and non-survivors having greater than 5.5 log_10_ PFU/mL (**Figure 4F**). Through the course of the study, all of the animals registered an elevated temperature between D4 and D8 PI, with treated survivors returning to baseline temperature by D10 PI (**Figure 4G**). Clinical scores and severe thrombocytopenia only appeared in non-survivors (**Figure 4H and 4I**). Thus, MBP431 can prevent onset of EBOV disease when administered as PEP and can resolve milder courses of EBOV disease when delivered IM at a 5 mg/kg IM dose.

**Figure 4.**
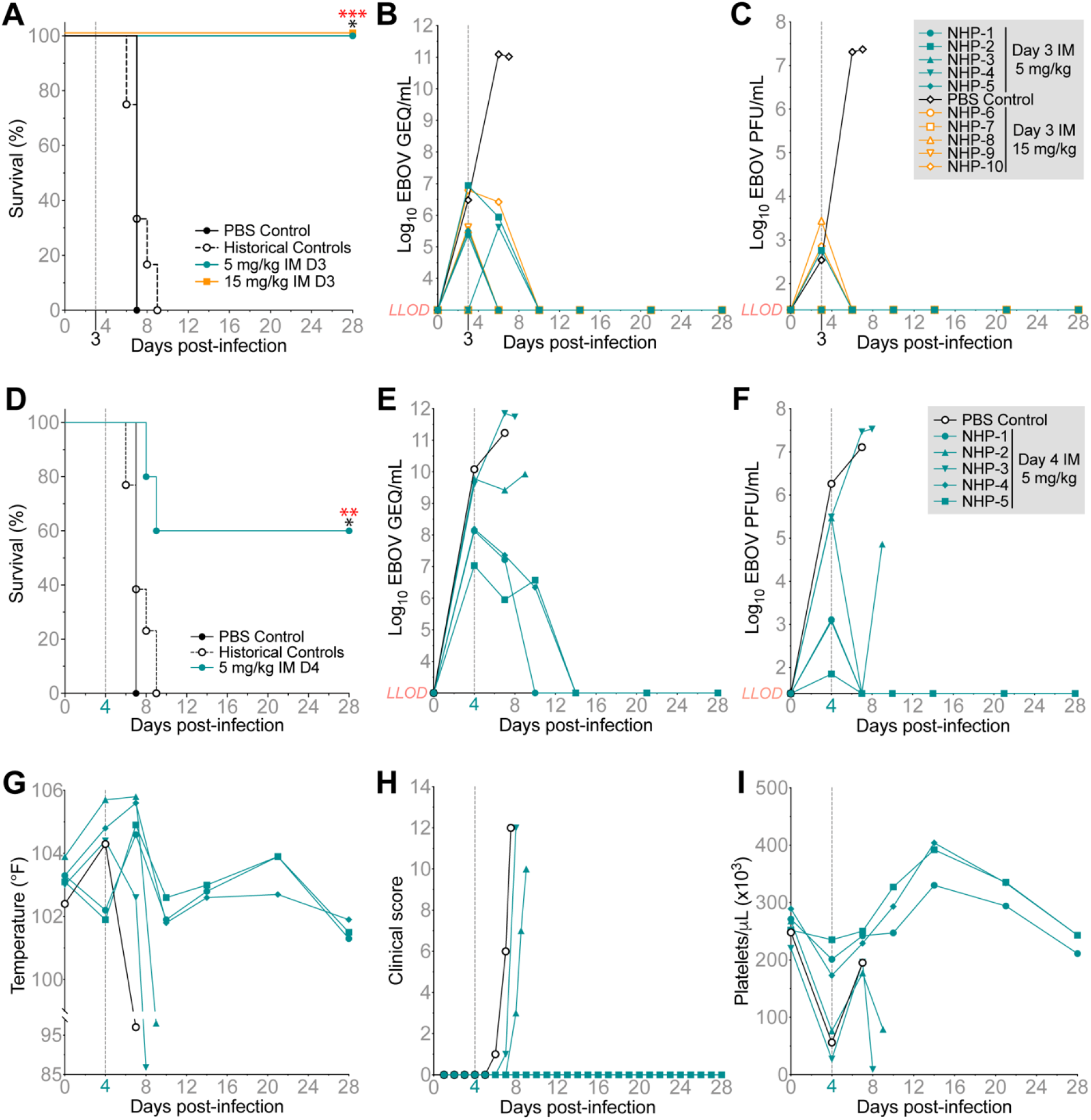
Evaluation of MBP431 as a PEP or therapeutic IM injectable treatment in NHPs challenged with EBOV/Kikwit. **(A)** Survival curves for NHPs challenged IM with 1000 PFU of EBOV/Kikwit. Animals received either a single 5 mg/kg (cyan) or 15 mg/kg (orange) IM dose of MBP431 on D3 PI. Historical controls and the in-study control are shown in black. Statistical significance was determined using a Log-rank Mantel-Cox test, against the in-study PBS control alone yielding a P value of 0.0253 (*) or combined with the 13 historical controls a P value of 0.0003 (***, red) for both treatment groups. **(B)** Average GEQ/mL of EBOV/Kikwit and **(C)** infectious EBOV/Kikwit (PFU/mL) in the blood of each NHP treated D3 PI. **(D)** Survival curves for NHPs challenged IM with 1000 PFU of EBOV/Kikwit with the 5 mg/kg IM dose of MBP431 walked out to D4 PI. Statistical significance was determined using a Log-rank Mantel-Cox test, against the in-study PBS control alone yielding a P value of 0.0253 (*) or combined with the 14 historical controls a P value of 0.0045 (**, red). **(E)** Average GEQ/mL of EBOV/Kikwit and **(F)** infectious EBOV/Kikwit (PFU/mL) in the blood of each NHP treated D4 PI. **(G)** Body temperatures, **(H)** clinical scores, and **(I)** platelet counts for each NHP treated D4 PI. The legend for panels **(B-C)** appears in top right-hand corner of panel **(C)** (grey box), and the legend for panels **(E-I)** appears in the top right-hand corner of panel **(F)** (grey box). *LLOD* = lower limit of detection (3 log_10_ GEQ/mL and 1.39 PFU/mL).

## 1.4 DISCUSSION

Both the unprecedented 2013-2016 West African EVD epidemic and the ongoing SARS-CoV-2 global pandemic have underscored the need for having readily available medical interventions to halt the spread of infectious diseases. For EVD, both in West Africa and in outbreaks since, the high infection rates experienced by medical staff severely compromised the capability of ETUs to treat their patients (Evans et al., 2015). These issues were further exacerbated by the time and personnel required for IV infusion of treatments in low resource settings. Indeed, 12 patients enrolled in the PALM clinical trial succumbed to EBOV infection prior to receiving any available treatments (Mulangu et al., 2019). A medical countermeasure (MCM) targeting all ebolaviruses that could be rapidly administered via an IM injection to instantly prevent or reverse the course of infection could alter response and treatment paradigms during an ebolavirus outbreak. We evaluated MBP134, a potent pan-ebolavirus therapeutic comprised of two broadly neutralizing human mAbs, ADI-15878 and ADI-23774, as an easily administered IM injectable MCM to treat and/or prevent ebolavirus infection.

To expand the clinical utility beyond therapy for potential use as a long-term prophylactic or PEP, we introduced serum half-life mutations into the Fc regions of both mAbs in the MBP134 cocktail. The success of the prophylactic ring vaccination strategy used in Guinea to control the 2013-2016 EBOV epidemic was linked to its ability to stop new infections for 30 days after the 10-14 day window in which individuals remained at risk until their immune system responded to the vaccine (Bornholdt and Bradfute, 2018; Henao-Restrepo et al., 2017). In contrast to vaccines, which require varying periods of time to generate protective immunity, a single-dose IM-delivered immunoprophylactic could offer instantaneous protection, immediately disrupting the chain of transmission during an outbreak. Of the two half-life mutation variants, YTE and LS, analyzed in NHP pharmacokinetic studies, we observed the YTE mutations present in the MBP431 cocktail conferred a longer half-life. While sequence differences in a mAb variable region can lead to variable serum half-lives in the presence of either the YTE or LS mutations, we cannot rule out the possibility that the particularly short half-life of ADI-23774^LS^ (∼7 days) may be due to the development of anti-drug antibodies that were not generated from an equivalent exposure to ADI-23774^YTE^. Ultimately, MBP431 was tested for PEP and therapy, demonstrating significant protective efficacy with a single 5 mg/kg IM dose. With this dosing regimen, MBP431 did not appear to impact NHPs displaying a more advanced state of disease at time of treatment. While higher doses of MBP134 administered via IV have already demonstrated complete protection (Bornholdt et al., 2019), the 5 mg/kg IM dosing of MBP431 tested here represents a potentially realistic single dose IM regimen for an average weight human. It remains to be determined if lower doses of MBP431 tested here in combination with small molecule-based treatments like Remdesivir could amplify protective efficacy via augmented biodistribution in more severe cases as was observed in recent combination studies for the treatment of marburgvirus infection (Cross et al., 2021). Future experiments will also focus on the prophylactic utility of MBP431 when administered IM in NHP models of ebolavirus infection.

## 1.5 ACKNOWLEDGMENTS

The authors would like to thank the UTMB Animal Resource Center for husbandry support of laboratory animals and Drs. Kevin Melody, Chad Mire, Abhishek Prasad and Courtney Woolsey for assistance with the animal studies. Opinions, interpretations, conclusions, and recommendations are those of the authors and are not necessarily endorsed by the University of Texas Medical Branch.

This study was supported by the Department of Health and Human Services, National Institutes of Health grant numbers U19AI109711 and U19AI142785 to TWG and UC7AI094660 for BSL-4 operations support of the Galveston National Laboratory. Additionally, the efforts from Mapp Biopharmaceutical, Inc. were supported by U19AI109711 and U19AI142785. The pharmacokinetics study performed at the Tulane National Primate Research Center was supported, in part, by Grant OD011104 (CJR) from the Office of Research Infrastructure Programs, Office of the Director, NIH and Department of Health and Human Services, National Institutes of Health grant U19AI109762.

---

**Figure S1.**
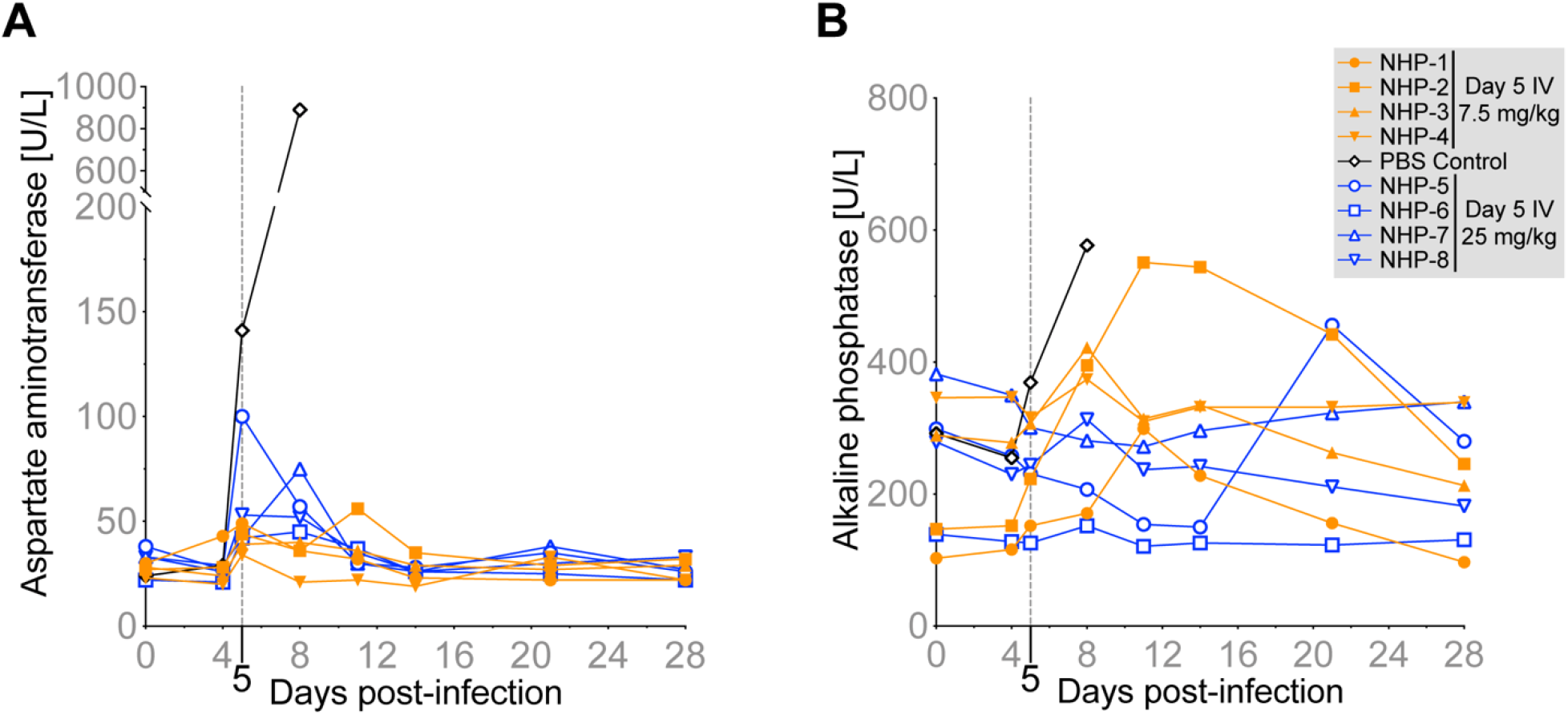
Graphical representation of aspartate aminotransferase (AST) and alkaline phosphatase (ALP) level from the therapeutic evaluation of MBP134 in NHPs challenged with SUDV/Gulu, Related to Figure 1. AST levels are graphed in panel **(A)** and ALP levels in panel **(B)**. The legend for the graphs in panel **(A)** and **(B)** appears in top right-hand corner of panel **(B)** (grey box).

**Figure S2.**
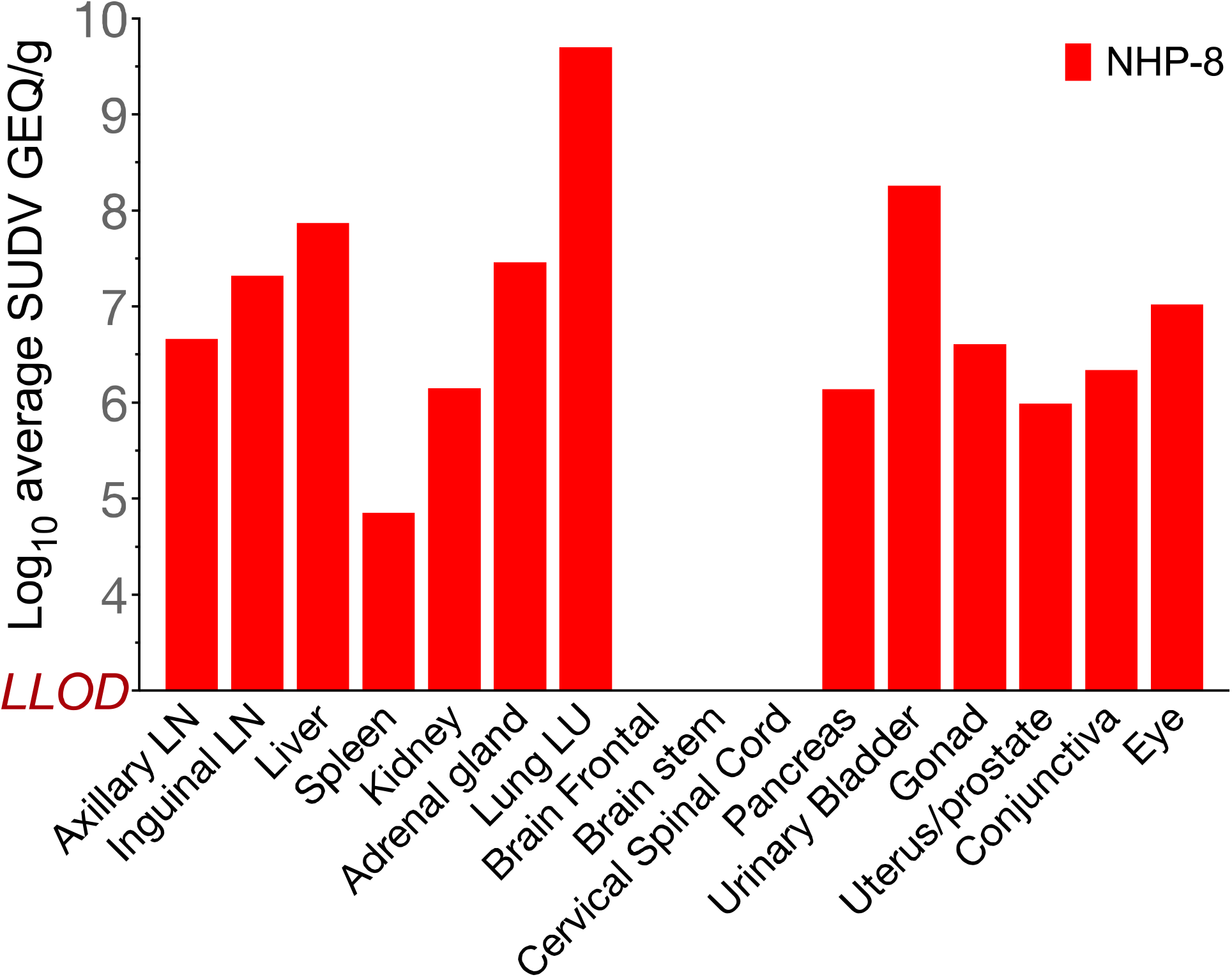
qRT-PCR of SUDV/Gulu genomic equivalents per gram (GEQ/g) present in tissues taken from NHP-8 challenged with SUDV/Gulu upon necropsy, Related to Figure 2. The viral load present in emulsified tissue samples at time of necropsy taken from NHP-8 is graphically displayed. NHP-8 had the highest viral load of all the Day 5 treated animals and the control animal on D5 PI. That viral load correlates with the severe systemic infection observed in the tissue samples taken from multiple organs in NHP-8.

